# Territorial risk of high pathogenicity avian influenza in France is associated with farmed species diversity

**DOI:** 10.64898/2026.10.10.758206

**Authors:** Clara Delecroix, Claire Guinat, Lisa Fourtune, Mathilde C. Paul, Timothée Vergne

## Abstract

Animal infectious diseases threaten food security, livestock production and public health. While disease prevention strategies typically target risk factors associated with high-competence hosts or environmental conditions, we hypothesise that lower-competence species can reduce disease risk by diluting effective contact rates between farms. Using spatial statistical models calibrated on three successive French high pathogenicity avian influenza (HPAI) epidemics (2020-2023), we show that greater local diversity of livestock farms, including cattle and pig farms, is associated with significantly lower HPAI risk. Moreover, we show that the positive association between duck houses and HPAI risk weakens as the number of less-competent non-duck poultry houses increases, revealing a dilution effect within poultry production systems. These findings suggest that livestock diversity is a previously overlooked determinant of disease spread and open new research avenues on territorial diversification to complement traditional approaches for the long-term control of high-impact animal infectious diseases.

## Introduction

Animal infectious diseases pose a significant threat to livestock production, farmers’ livelihood and food security, causing annual losses of billions of dollars globally (World Organization for Animal Health, 2026). They are also a major threat to public health with most emerging human diseases being zoonotic (Jones et al., 2008). Identifying disease risk factors helps mitigate disease impact and reduce the risk of future epidemics. These factors are usually identified by modelling the risk of disease occurrence as a function of putative risk variables, with such analyses often relying exclusively on variables related to high-competence species, i.e. species that can readily become infected and transmit the pathogen. For instance, the main investigated drivers of high pathogenicity avian influenza (HPAI) risk are related to poultry and wild bird distributions (Gilbert and Pfeiffer, 2012). Similarly, risk factors for foot-and-mouth disease (FMD) focus on the distribution of farms rearing ruminants and swine, the main competent host species for the virus (Kedang et al., 2025), while those for African Swine Fever (ASF) primarily involve pig- and wild boar-related factors (Boklund et al., 2024). Therefore, the potential influence of farmed species diversity, including low-competence or even non-host species, on animal infectious disease risk remains overlooked.

Previous experimental and modelling studies on several pathogens circulating in wildlife populations have shown that ecosystems with high species diversity are less suitable for pathogen circulation (Keesing et al., 2006; Keesing and Ostfeld, 2021; Ostfeld and Keesing, 2012). This pattern has been explained by a “dilution effect” in which low-competence hosts, i.e. hosts with reduced susceptibility and infectivity to a given pathogen, can limit pathogen spread by reducing effective contact rates between high-competence species (Keesing and Ostfeld, 2021). This mechanism has mostly been documented for vector-borne diseases, where low-competence hosts divert vector bites away from competent hosts, reducing the proportion of infectious vector bites available for onward transmission. For instance, deer species composition was shown to influence the risk of outbreak emergence for several tick-borne pathogens (Fabri et al., 2024), and host composition could reduce the transmission of West Nile virus (Marini et al., 2017; Martins et al., 2022). Beyond vector-borne systems, a dilution effect has also been suggested in non-vectorial, directly transmitted systems, arising through contact-based transmission pathways when species or production units differ in their epidemiological competence. For example, exposing amphibian species with varying competence to the trematode *Ribeiroia ondatrae* reduced the number of infected individuals in an experimental study (Johnson et al., 2013). For livestock diseases specifically, such a dilution effect has also been observed: in cattle, the presence of bovine tuberculosis was found to decrease with increasing mammal species richness (Huang et al., 2013). Building on this evidence, we explore whether a similar dilution effect may operate more broadly for between-farm transmission of livestock diseases: by increasing the heterogeneity of production systems, greater farmed species could reduce indirect effective contacts between farms rearing competent host species, for example those mediated through rendering trucks, veterinary visits, or shared equipment (Figure 1A and 1B).

**Figure 1.**
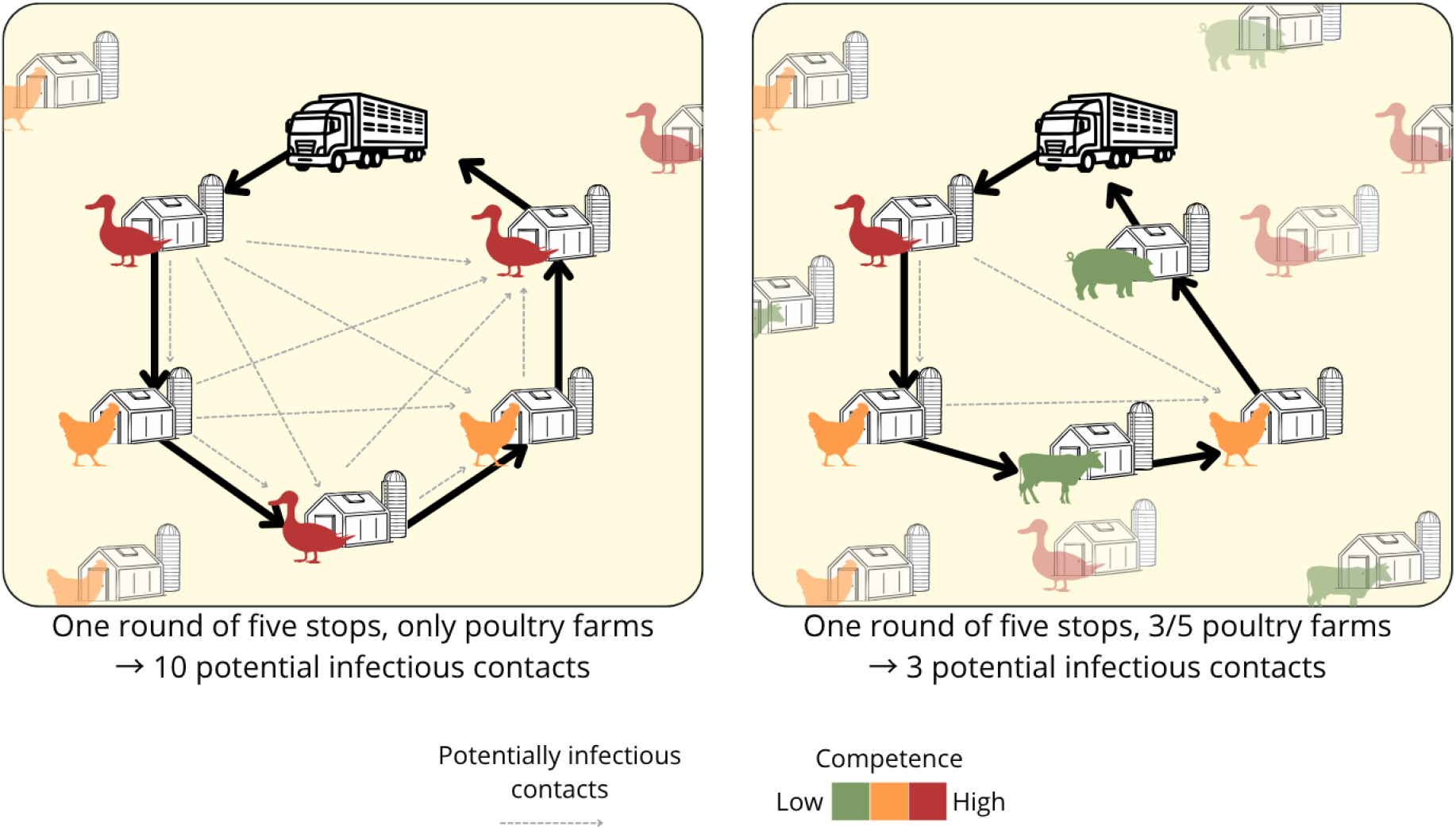
Conceptual scheme of how a dilution effect may operate for livestock diseases. A: transmission in a diversity-poor context, all species are highly competent and thus highly connected via indirect effective contacts. B: transmission in a highly diverse context, the density of highly competent species is the same but the effective contact rate decreases as the contacts are distributed among species.

This question actually arises for HPAI, which remains a global threat to animal and human health (Koopmans et al., 2024) and whose host range has broadened in recent years: beyond wild birds and poultry, sporadic to widespread infections have been detected in other livestock species, such as cattle and pigs (Long et al., 2019; Peacock et al., 2024). The epidemiological contribution of these newly affected species in disease occurrence remains unclear. Yet, previous work on species diversity and HPAI risk has been restricted to comparisons within wild bird communities, where the composition and diversity of wild bird species have been linked to variation in avian influenza prevalence and spread (Huang et al., 2019; Yin et al., 2024). Diversity across species of markedly different competence, including non-avian and non-host species, has not been considered, despite its potential to shape the occurence risk.

France constitutes a particularly relevant setting to investigate this question, given the diversity of its livestock production systems and being one of the most affected European countries by HPAI. Poultry farms, in particular duck farms, have been heavily aftected, along with cases in wild birds and sporadic cases in wild mammals (Lambert et al., 2022; Scoizec et al., 2024). No cattle outbreaks have been reported, and serological surveys conducted in dairy cattle following the recent epidemics have so far shown no evidence of previous exposure (Buczkowski et al., 2026). Previous studies have shown that, although poultry farm outbreaks were initially seeded by incursions from wild birds, the very high number of infected poultry farms has been driven primarily by farm-to-farm transmission (Briand et al., 2022; Guinat et al., 2025). These studies have also shown the role of ducks as the most susceptible species, whose high density has contributed to the large number of outbreaks (Andronico et al., 2019; Bauzile et al., 2023; Guinat et al., 2019a). However, the role of chickens and other non-duck poultry species has yielded contrasting findings, sometimes positively, sometimes negatively associated with risk (Gkrinia et al., 2025; Guinat et al., 2019b), which could be due to the different competence compared with duck species (Filaire et al., 2022). Thus, the role of chicken species may depend on farmed species composition, diluting contacts and thus reducing transmission risk when species with a higher competence are present while having the opposite effect in the presence of lower competence species. In addition, while much attention has focused on poultry, the potential influence of other farmed species has received little consideration. Indeed, across the three most recent French epidemics, although most municipalities affected by HPAI had very high duck farm densities, they were also highly specialised, with an average of 1.8 farmed species per municipality, compared with 3.8 species in municipalities where no outbreaks were reported (Supplementary Information). As a proof of concept, we investigated indirect contact patterns between poultry farms via rendering trucks and found that territorial diversity of farmed species was negatively associated with the number of poultry farms visited in one round, reducing the number of potentially infectious contacts (Supplementary Information). This pattern raises the question of whether livestock diversity acts as an overlooked ecological determinant of HPAI transmission, and remains to be formally assessed.

In this study, we hypothesised that the dilution effect mediated by the diversity of farmed species introduced above, reduced indirect effective contact rates between high-competence duck farms in territories with greater livestock density, thereby influencing HPAI transmission. To test this, we examined risk factors associated with spatial distribution of HPAI outbreaks across three epidemics that occurred in France between 2020 and 2023, focusing primarily on variables related to low-competence farmed species. Using one spatial statistical model per epidemic, we quantified outbreak risk as a function of a range of explanatory variables. If supported, the dilution effect could open new research avenues on how to improve the resilience of livestock systems across Europe to HPAI outbreaks.

## Results

### Modelling HPAI outbreak occurrence

Three large HPAI epidemics affected France in 2020-2021, 2021-2022 and 2022-2023, with respectively 451, 1323 and 795 recorded outbreaks in poultry farms (Figure 2). While the first one was restricted to the South-West of the country, highly specialised in duck production, the next two epidemics recorded outbreaks in the whole country. In addition, 839 cases were recorded in wild birds over the period of interest, whose spatial distribution was biased to areas subject to surveillance and showed a minor role compared to farm-to-farm transmission, and are thus excluded from subsequent analyses (Briand et al., 2022; Guinat et al., 2025). Three spatial statistical models were fitted to the domestic poultry outbreaks of three epidemics of HPAI in France using R-INLA (Moraga, 2019). For each epidemic, the models used the number of outbreaks within a spatial unit (4-kilometre-wide hexagons) as the outcome variable, and included spatial and non-spatial random effects within a Poisson modelling framework. The effect of four explanatory variables was tested: the number of duck houses per hexagon, ranging between 0 and 83, the number of non-duck poultry houses, including in majority chicken and turkey, ranging between 0 and 113, the proportion of low-competence species in the farms, defined as the proportion of non-poultry farms (pig, cow, sheep and goat) across all farms present in the hexagon, as well as the proportion of the area covered by water.

**Figure 2.**
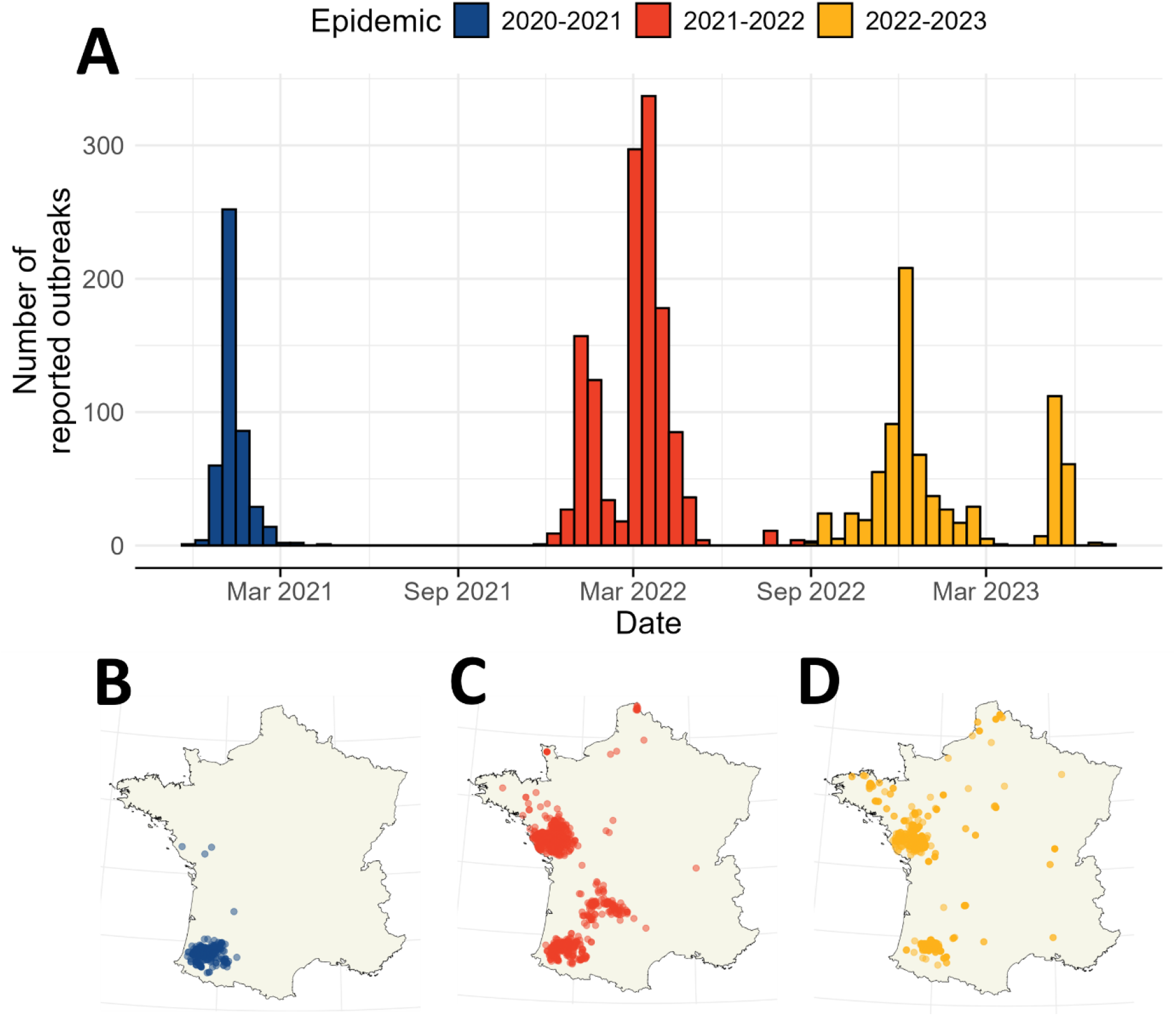
Outbreak data in domestic poultry during three HPAI epidemics in France between September 2020 and 2023. A: Temporal distribution of the outbreaks. B: Spatial distribution of the outbreaks in 2020-2021. C: Spatial distribution of the outbreaks in 2021-2022. D Spatial distribution of the outbreaks in 2022-2023.

### Predictors of HPAI outbreak occurrence

Four explanatory variables were selected as significantly improving the model for at least one epidemic, using a forward selection procedure. Figure 3 illustrates the effects of the predictive variables across the three epidemics and suggests that, even if the effects were not systematically statistically significant, they were consistent between epidemics. Overall, while the number of duck houses and non-duck poultry houses was positively associated with HPAI risk (for zero non-duck poultry houses and zero duck houses, respectively, as an interaction was also included). The proportion of low-competence farmed species and the proportion of surface covered by water was negatively associated with HPAI risk, regardless of the other variables. The hydrology variable was considered as a proxy for water wild bird species, involved in HPAI transmission (Couty et al., 2026), and the protective effect was unexpected. This suggests that either wild bird species did not play a significant role in the three studied epidemics, or the proxy was inappropriate.

**Figure 3.**
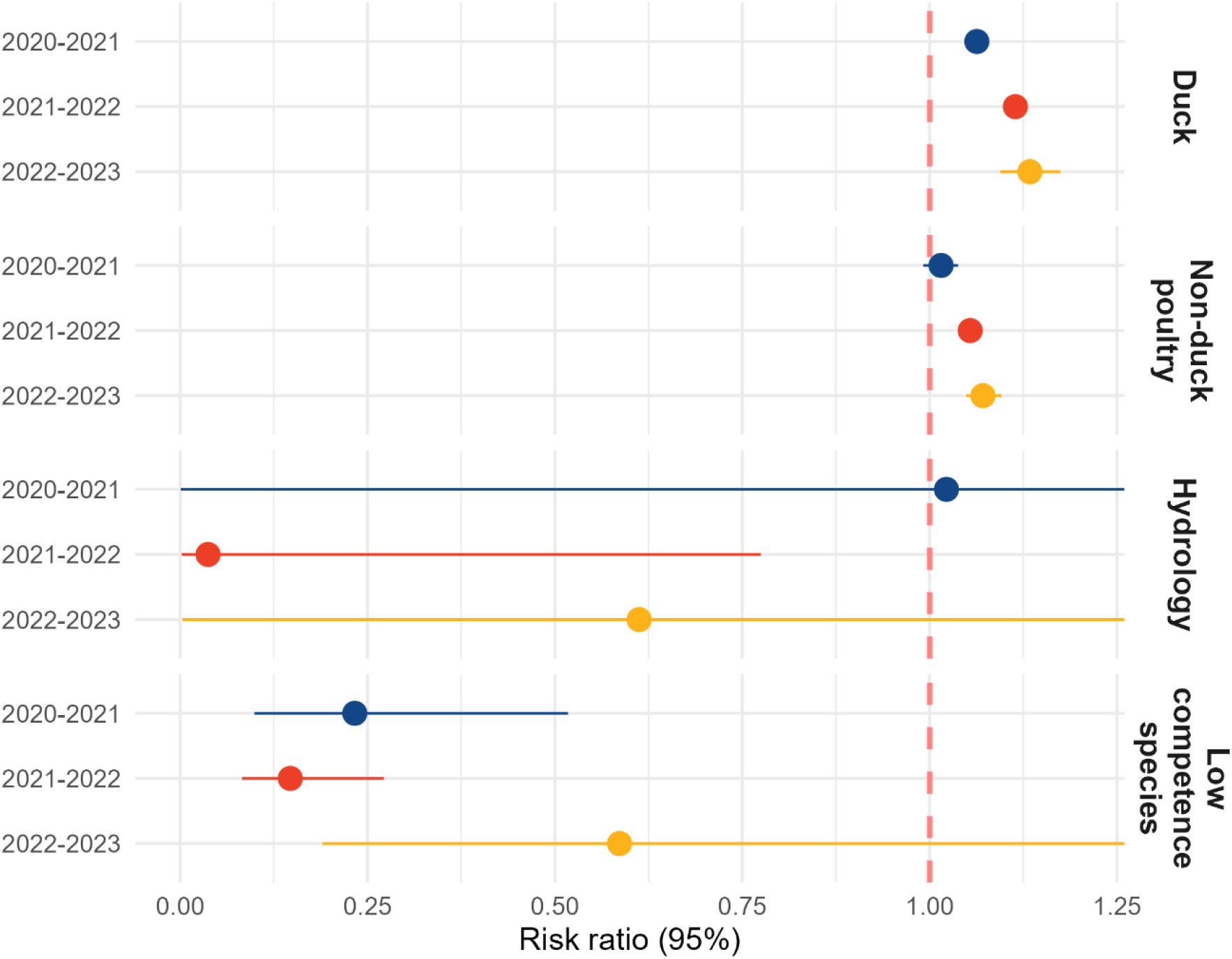
Median risk ratio and 95% credible intervals of the coefficients of the predictive variables in the different models. The coefficient for the variable *duck* should be interpreted at zero *non-duck poultry* houses, as there is an interaction between *duck* and *non-duck poultry*. Similarly, *non-duck poultry* should be interpreted at zero *duck* houses. Hydrology refers to the proportion of surface area covered by water and low competence species refers to the density of farms rearing either cattle or pigs.

### Indication of a dilution effect

The proportion of low-competence farmed species showed a consistent negative association with HPAI risk across all three models, reaching statistical significance for two of them (Figure 3), with risk ratios of 0.23 [0.09, 0.52], 0.15 [0.08, 0.27] and 0.59 [0.19, 1.93], respectively (Table S2). Thus, for a given value of duck and non-duck poultry house densities, a greater proportion of low-competence species was associated with a lower risk of outbreak occurrence.

Additionally, including an interaction term between duck and non-duck poultry house densities also improved two of the three models (evaluated using the WAIC), suggesting that the association between each type of poultry house and HPAI depended on the number of the other. As expected, increasing the number of duck houses within a unit increased the probability of observing at least one outbreak, but this increase was attenuated in areas with higher numbers of non-duck poultry houses (Figure 4A and C). Similarly, the association between the number of non-duck poultry houses and HPAI risk varied depending on the level of ducks: at low duck house numbers, increasing the number of non-duck poultry houses was associated with increased HPAI risk, but this association weakened as duck house numbers increased and even became negative at very high duck house densities (Figure 4A and B).

**Figure 4.**
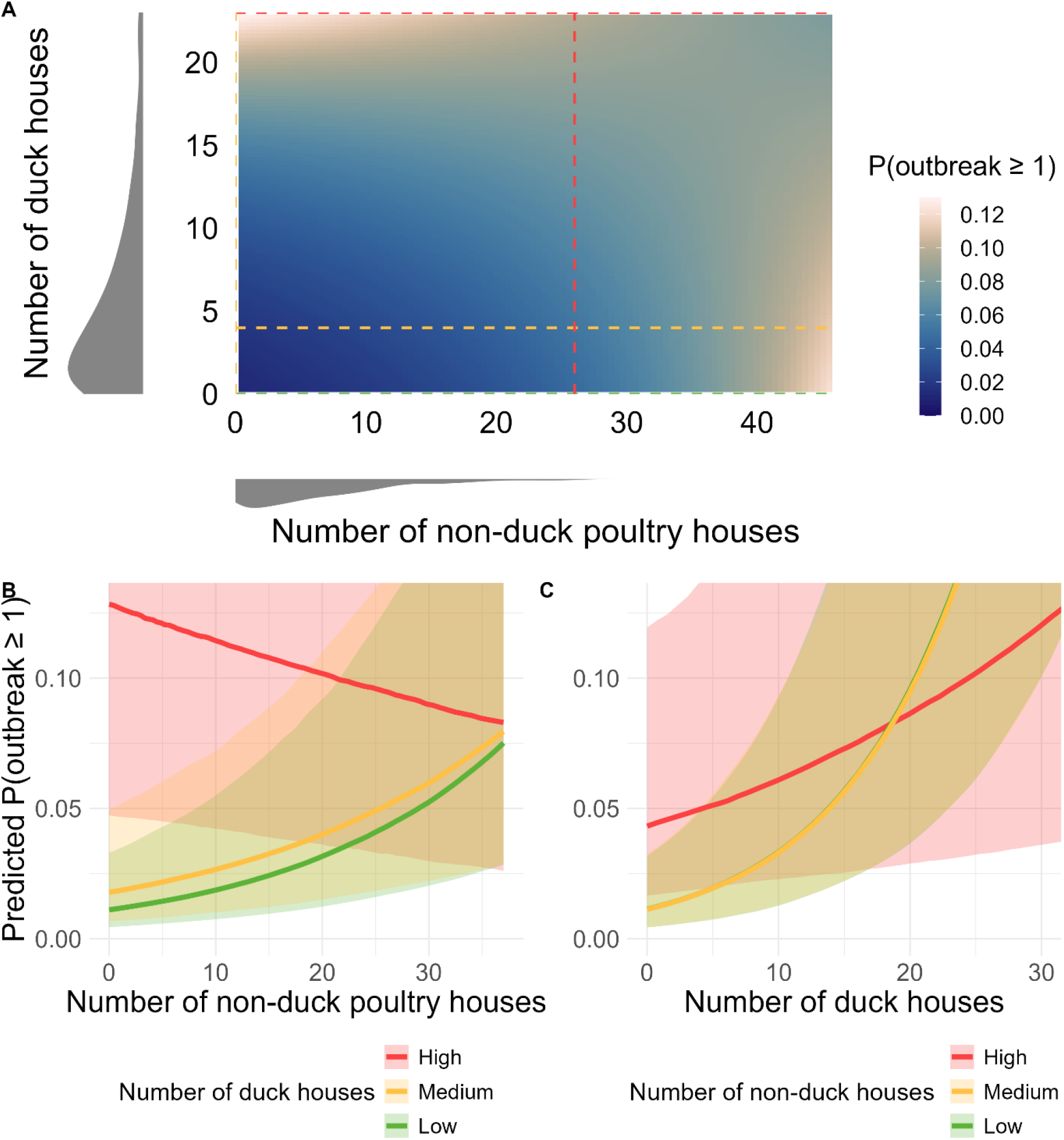
A: Interaction effect between the number of duck and non-duck poultry houses, for the model 2021-2022 (consistent with other models, provided in SI) . The probability of having at least one reported outbreak in the spatial unit as a function of the number of duck houses and number of non-duck poultry houses, calculated for other variables at their average values, without spatial component. The three levels of figure B are represented as horizontal lines, and the three levels of figure C as vertical lines. B: Probability of having at least one reported outbreak in the spatial unit as a function of other poultry house number, for three levels of numbers of duck houses (2.5th, 50th and 97.5th percentile), other variables set at their average values, calculated without spatial component. C: Probability of having at least one reported outbreak in the spatial unit as a function of duck house number, for three levels of numbers of non-duck poultry houses (2.5th, 50th and 97.5th percentile), other variables set at their average values, calculated without spatial component.

Together, these results suggest the existence of a dilution mechanism, whereby landscapes with greater diversity of farmed species with different levels of competence are associated with reduced HPAI risk. This mechanism may result from reduced effective contact rates between poultry farms, either through a greater presence of farms raising low-competence non-poultry species, such as cattle and pigs, or through increased heterogeneity within poultry production systems due to the presence of less-competent non-duck poultry farms.

## Discussion

To investigate whether the dilution effect hypothesis applies the context of farmed animals, we used spatial risk models of outbreak occurrence to quantify the association between geographical HPAI risk and the distribution of livestock farms across species with varying levels of competence for HPAI. Our results provide preliminary evidence supporting this dilution mechanism through two complementary patterns. First, the proportion of farms rearing low-competence species was negatively associated with the HPAI outbreak risk for two of the three epidemics. Second, a statistically significant negative interaction between duck and non-duck poultry house densities suggested that the positive association between the density of one poultry type and HPAI risk was weaker in areas with higher densities of the other poultry type. These findings were robust across three models, one per HPAI epidemic in France since 2020.

We showed that an increased proportion of low-competent species was significantly associated with a reduced risk of HPAI outbreaks in two of the three epidemics, regardless of duck and non-duck poultry house densities. This association may result from a broad dilution of indirect contacts between poultry farms, particularly those mediated through non-species-specific routes such as rendering truck rounds (Supplementary Information). Because rendering trucks are not dedicated to any particular species, indirect effective contact rates through these routes are likely to depend on local livestock diversity, meaning that contact opportunities between farms rearing competent species may vary according to the composition of surrounding farmed species. Although our proof of concept of diversity-dependent contact rates focused on rendering truck rounds, other mechanisms may generate similar patterns. For example, sharing equipment between farms could generate diversity-dependent effective contact rates if such exchanges occur between farms rearing different species. Similarly, veterinary visits could contribute to diversity-dependent contact patterns if veterinarians in more diversified regions interact with a broader range of farm types. This concept of diversity-dependent effective contact rates opens new avenues for characterising contact patterns among farms and improving our understanding of infectious disease transmission dynamics.

Furthermore, a significant interaction was found between duck and non-duck poultry house numbers, as the direction of their respective effects depended on the value of the other. Increasing non-duck poultry house numbers was associated with higher outbreak risk when duck house numbers were low (2.5% quantile), but with lower outbreak risk when duck house numbers were high (97.5% quantile). Similarly, the increase in risk associated with duck house numbers was attenuated in areas with higher densities of non-duck poultry houses. A possible explanation for this pattern might be the reduced susceptibility and infectivity of non-duck poultry farms in France compared to ducks (Andronico et al., 2019). At high duck house densities, increasing the numbers of non-duck poultry houses with lower competence may dilute contact opportunities between duck houses and reduce transmission risk. Consistent across the three epidemics of HPAI in France, these findings suggest a dilution mechanism whereby reducing territorial specialisation in farmed species, even within poultry production systems, may decrease contact rates between highly competent species and enhance the robustness of poultry systems to future HPAI outbreaks.

The dilution effect has sparked many debates in eco-epidemiology (Randolph and Dobson, 2012; Rohr et al., 2020). Previous research (Ostfeld and Keesing, 2012) established three necessary conditions for a dilution effect to occur: (i) hosts differ in their competence for pathogen transmission, (ii) highly competent hosts tend to occur in contexts with low diversity while less competent hosts are found in diverse contexts and (iii) the presence of less competent hosts reduces the abundance or (indirect) contact rates between highly competent hosts. Condition (iii) is often observed for vector-borne diseases, where diverse communities dilute vector feeding and thereby reduce exposure of competent hosts. In livestock systems, a dilution effect is also expected to operate only in specific conditions. First, consistent with condition (i), farmed species must differ substantially in their competence. If all farmed animals have the same competence, no dilution can occur. In the context of HPAI, non-poultry species can be infected but appear to contribute little to onward transmission in France (Flavia et al., 2024), whereas ducks are the main drivers of transmission and non-duck poultry species contribute to a lesser extent (Guinat et al., 2019b). Second, consistent with condition (iii), a substantial proportion of between-farm contacts must be influenced by local farmed species diversity. If most contact rates between competent farms are species-specific, farm diversity will not alter transmission opportunities. Beyond HPAI, similar dilution mechanisms may apply to other livestock diseases involving different hosts with different competence in transmitting the disease, such as FMD, where cattle and pigs are highly competent hosts and small ruminants have lower competence (Barnett and Cox, 1999), or ASF, where pigs are the primary host and other livestock species cannot be infected.

Further research is necessary to better characterise the mechanisms underlying this potential dilution effect on HPAI. First, our analysis simplified the poultry system into a duck population and a non-duck poultry population based on the known differences in susceptibility and infectivity between these farm types. However, poultry systems within species are themselves more heterogeneous than this, with a diversity of production systems, quality labels and of biosecurity practices that are known to be associated with virus exposure and, therefore, with HPAI risk (Delpont et al., 2018; Hautefeuille et al., 2026). In future work, it will therefore be important to account not only for poultry species but also for farm types, which could contribute to strengthening or weakening the association found in this study. Second, it remains unclear how changes in farmed species diversity alter between-farm contact patterns and which types of contacts contribute to this phenomenon. Addressing these questions will require characterising between-farm contact networks and distinguishing species-specific contacts from non-species-specific contacts that may mediate the dilution effect. Such characterisation could help calibrate mechanistic models more realistically, and improve our understanding of how this mechanism operates. Ultimately, combining compartmental epidemiological models with realistic between-farm contact data could help identify the conditions under which dilution mechanisms emerge and persist.

Increased diversity of farmed species in territories might contribute to mitigate the risk of livestock disease occurrence in specific epidemiological contexts. However, such diversification may also increase opportunities for pathogen exposure among low-competence species, raising questions about the potential for host adaptation. For instance, the host range of HPAI is constantly evolving, and further research is necessary to understand how increasing farm diversity may influence the risk of viral adaptation to new hosts. An increasing number of HPAI spillover events into mammalian species, including swine and cattle, have been reported in recent years (Bourret, 2018; Neumann and Kawaoka, 2024). In the present study, these species are categorised as low competence because limited transmission among mammals has been documented and their seroprevalence remains low in Europe (Flavia et al., 2024). However, a greater diversity of production systems in territories could increase the frequency of contacts between poultry farms and other livestock farms, potentially creating additional opportunities for viral adaptation and spillover (Rohr et al., 2020). Conversely, our findings suggest that such diversification may reduce overall transmission intensity through a dilution mechanism, thereby potentially limiting the number of successful transmission chains available for viral evolution. Further investigations are needed to determine how to harness the benefits of the dilution mechanism while minimising the risk of pathogen adaptation to new host species. This study is only a first step in investigating a dilution effect affecting livestock disease transmission. Future work should investigate the generalisability of this theory with other pathogens and in other contexts.

## Conclusion

Using HPAI as a case-study, our study provides initial evidence suggesting a dilution mechanism in animal infectious disease systems. This initial evidence opens new research avenues on the association between reduced territorial specialisation and infectious diseases risk. Moving forward, confirming these initial findings in other contexts and for other pathogens and characterizing their potential side effects would contribute to finding leverages to increase the robustness of our production systems to infectious disease threats. In the long term, these leverages could highlight structural changes in the spatial organisation of livestock production systems as an alternative for reactive responses to emerging disease events, as is currently common in Europe.

## Materials and methods

### Data

The country was divided into 40,807 4-kilometre-wide hexagons, corresponding to the average size of a French municipality. The hexagons served as the spatial unit of the model, and the risk was subsequently analysed for each hexagon.

Data related to confirmed HPAI poultry outbreaks between September 2020 and August 2023 were obtained from the French Ministry of Agriculture (DGAL). An outbreak was defined as a poultry house with at least one PCR-confirmed infected bird. A poultry house was defined as a building containing poultry flocks, belonging to a farm and identified by a unique reference code, called INUAV (unique identifier of poultry buildings). Three epidemics were considered (2020-2021, 2021-2022, 2022-2023), each spanning the period from 1 September of a given year to 31 August of the following year (Figure 2). The number of outbreaks in each spatial unit was used as the response variable.

Data on the poultry house population, including their establishment dates and closing dates, i.e when they were administratively registered as a poultry house, were also obtained from the DGAL. Active poultry houses for each year were identified as houses with an establishement date before the end of the considered time period, and a closing date after the beginning of the considered time period. Poultry houses were divided into duck houses (containing at least one duck) and non-duck poultry houses. The numbers of active duck houses and of active non-duck poultry houses were calculated in each spatial unit for each epidemic period and used as predictors.

To explore a potential dilution effect, data on other farming systems were extracted from the publicly available Agreste database (Agreste, 2022) which contains a census of cattle, sheep, goat, pig, duck and chicken farms, conducted in France in 2020. This census was used to calculate the proportion of low-competence hosts in each spatial unit, as the proportion of non-poultry species farms, i.e species that have a low competence in transmitting HPAI, among all farms, which was used as a predictor.

Finally, the proportion of surface covered by water, used as a proxy for the presence of wild water birds (Gilbert and Pfeiffer, 2012), was extracted from the IGN database (Institut national de l’information géographique et forestière (IGN), 2025) and also used as a predictor.

For each epidemic, the study area used to train the models was defined as the concatenation of all hexagons where at least one outbreak was reported during that epidemic and of all those located within 10 km from reported outbreaks.

### Model

For each epidemic year independently (2020-2021, 2021-2022, 2022-2023), spatial generalised mixed Poisson models were used to quantify the effect of the predictive variables on the number of outbreaks reported in each hexagon of the study area. To account for spatially and non-spatially structured effects, a Besag-York-Mollie (BYM) conditional autoregressive (CAR) modelling structure was used (Besag et al., 1991; Blangiardo et al., 2013). A BYM structure includes both an intrinsic CAR effect based on the first-order adjacency between hexagons, and an independent and identically distributed (iid) Gaussian random effect. Models were fitted for each epidemic using the integrated nested Laplace approximation (INLA) in R, using the package R-INLA (Moraga, 2019).

### Variable selection

First, correlation between the variables was checked, but no variable was removed as no Pearson correlation coefficient was above 0.6. Then, predictive variables were selected via a forward selection procedure, using the widely applicable information criterion (WAIC), which assesses the out-of-sample predictive performance. Variables were added sequentially until the WAIC could not be improved by at least 5 points. The forward selection procedure was run for each epidemic year independently, and the variables selected for at least one epidemic year were kept for the final model to maintain coherence across the three models.

Finally, we tested whether an interaction between duck and non-duck poultry house densities improved model performance, as would be expected under the dilution effect hypothesis. Moreover, this interaction could help explain previously inconsistent findings regarding the relationship between non-duck poultry species and HPAI risk (Gkrinia et al., 2025; Guinat et al., 2019). For parsimony reasons, additional interactions between other variables were not tested.

## Acknowledgements

The authors thank the General directorate for Food of the French Ministry of Agriculture, Food processing and Food Sovereignty (MAASA) for providing the data. We are grateful to the INRAE MIGALE bioinformatics facility (MIGALE, INRAE, 2020, doi: 10.15454/1.5572390655343293E12) for providing help and computing and storage resources.

