## Supplementary information for "Territorial risk of high pathogenicity avian influenza in France is associated with farmed species diversity"

Belonging to the manuscript titled "**Territorial risk of high pathogenicity avian influenza in France is associated with farmed species diversity.**"

Clara Delecroix, Claire Guinat, Lisa Fourtune, Mathilde C. Paul and Timothée Vergne

### Description of the production types in municipalities in France

For each municipality, we summarised the number of outbreaks that occurred there and compared municipalities where outbreaks were reported (n=730) against municipalities with no reported outbreak (n=34 238). Each municipality's production is assessed using the Agreste database (Agreste, 2022). The Agreste database is a census of the type of farms present in the municipalities, considering the number of cows, sheep, goat, pig, chicken and duck farms as the different species. Difference in duck or chicken species for instance are not accounted for in the analyses. We assess the average number of distinct farmed species across municipalities in both groups.

### Evidence of a diversity-dependent contact rate between poultry farms via rounds of rendering truck

We evaluated the effect of territorial diversification of farmed species on between-poultry-farm contact rates associated with rendering truck rounds. Data from a French rendering company that collects animal carcasses from farms and operates in the southwestern quarter of France were obtained for the period from September 2016 to December 2018. The data detail truck rounds, with a round being defined as a single truck visiting a sequence of farms on a given day, and contain information on the truck identification, the date of the round and the characteristics of the farms visited during that round (including location and species collected). For each round, we extracted the round length, defined as the number of farms visited, the number of poultry farms visited, as well as the municipalities visited. For simplicity, we assumed that all farms visited within the same round were in contact, regardless of the order of visits. To characterise the production context of the area where a round took place, we computed the average proportion of low-competent species farms in the visited municipalities. This proportion was computed using the publicly available Agreste database (Agreste, 2022) which contains a census of cattle, sheep, goat, pig, duck and chicken farms, conducted in France in 2020. The proportion of non-poultry farms in each visited municipality was calculated as the proportion of non-poultry farms among all farms.

Using a Poisson regression, we quantified the association between the proportion of low-competence hosts (predictor variable) and the number of poultry farms visited during a round (response variable). We considered the number of poultry farms visited during a round as a proxy for the effective contact rate associated with rendering trucks. To ensure that this relationship held regardless of the number of poultry farms in the visited municipalities, we fitted a second model using the number of poultry farms as a categorical predictor, with categories defined based on the quartiles of the number of poultry farms (0-25%: low; 25-50%: medium-low; 50-75%: medium-high; 75-100%: high). For both models, increasing the proportion of low-competence farms was statistically significantly and negatively associated with the number of poultry farms visited during a round. These results show that the number of indirect contacts between poultry farms associated with rendering truck rounds is reduced in areas with a higher diversity of low-competence species, irrespective of the number of poultry farms.

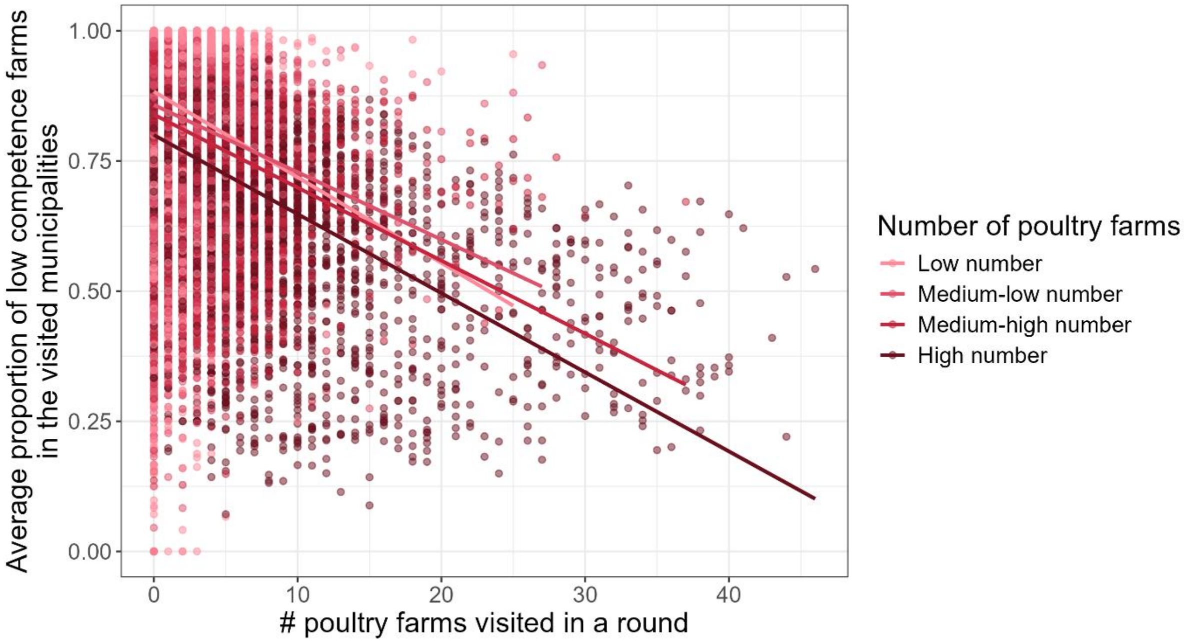


Figure S1: Number of poultry farms visited in a round of a rendering truck, as a function of the average proportion of low-competence species farms in the visited municipalities. Linear regression is calculated for each group of the number of poultry farms in the visited municipalities, grouped into quartiles (0-25%, 25-50%, 50-75%, 75-100%]).

Table S1. Risk ratio of the proportion of low-competence species to explain the number of poultry farms visited in a round of a rendering truck and its associated p-value. Model 1 includes the proportion of low-competence species as the only predictor, and Model 2 includes the number of poultry farms in the visited municipalities as an additional predictor.

| Model | Risk ratio of the variable “proportion of low competence species” | Associated p-value |
|---|---|---|
| Model 1 | 0.041 | <2e-16 *** |
| Model 2 | 0.064 | <2e-16 *** |

### Predictive variables used in the model

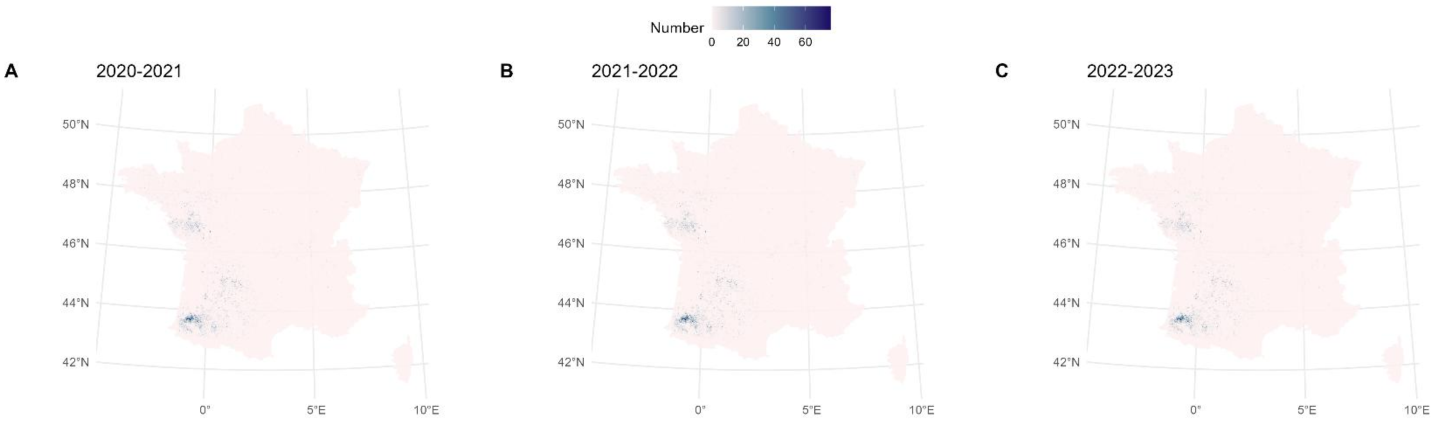


Figure S2: Number of active (defined as administratively open before September 1st and closed after August 31st of the year of interest) duck houses per hexagon. A Active duck houses for the epidemic year 2020-2021 B Active duck houses for the epidemic year 2021-2022 C Active duck houses for the epidemic year 2022-2023

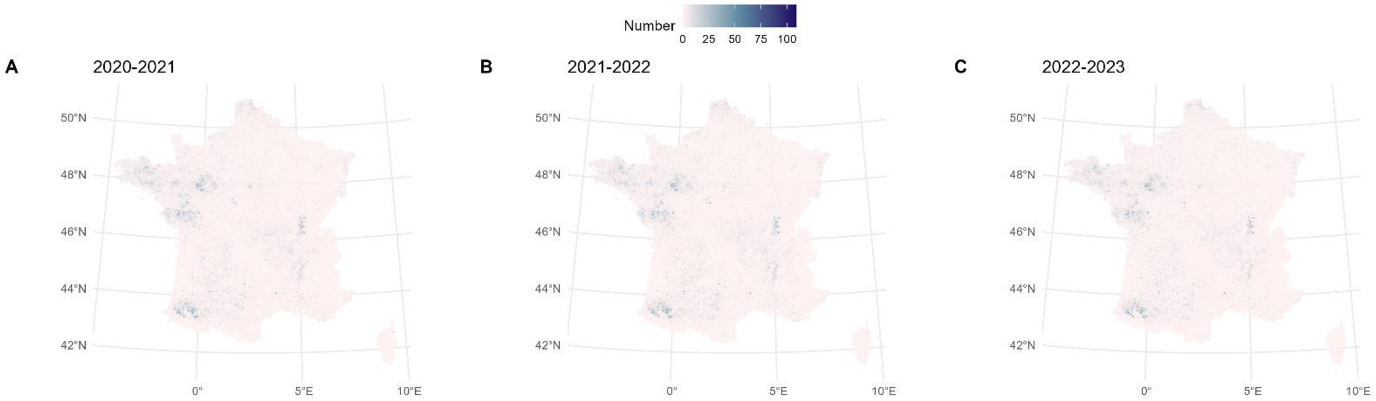


Figure S3: Number of active (defined as administratively open before September 1st and closed after August 31st of the year of interest) non-duck poultry houses per hexagon. A Active non-duck poultry houses for the epidemic year 2020-2021 B Active non-duck poultry houses for the epidemic year 2021-2022 C Active duck houses for the epidemic year 2022-2023

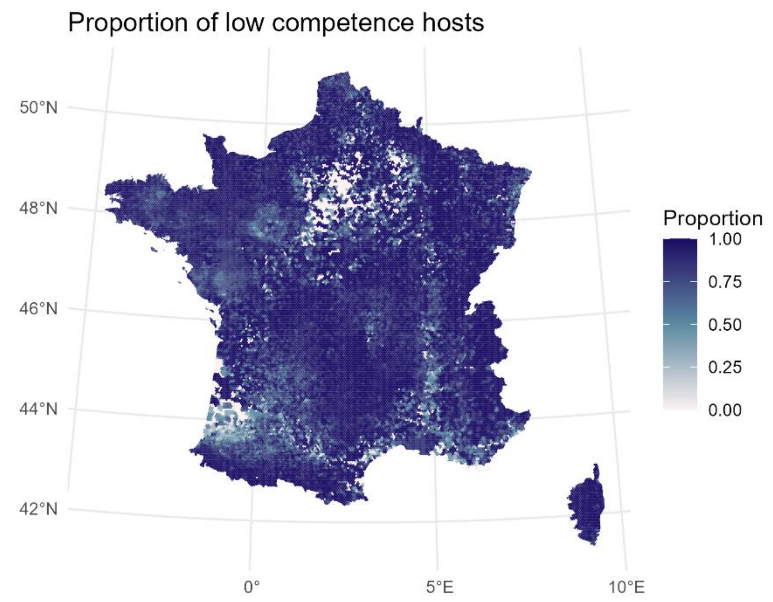


Figure S4: Proportion of low-competence species, defined as non-poultry farms (cows, pigs, sheep, goat) per hexagon

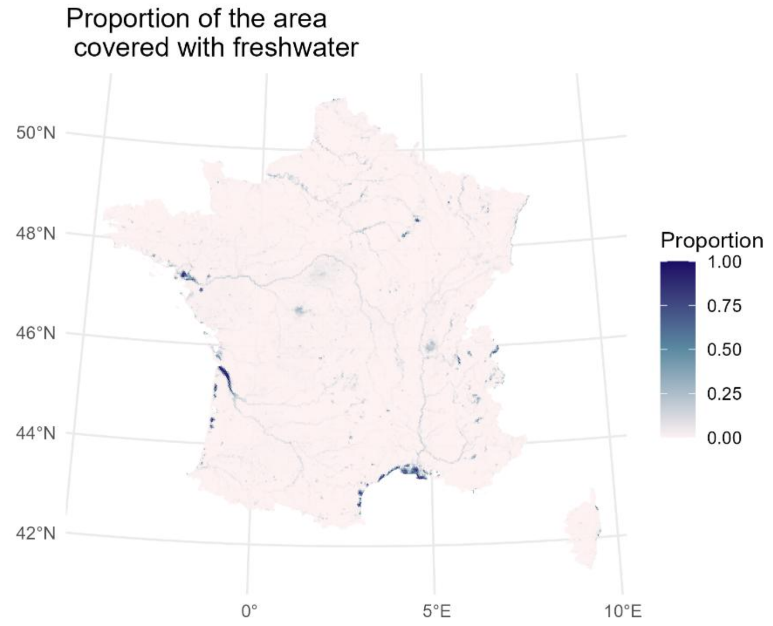


Figure S5: Proportion of the surface of the hexagone covered by water

### Risk ratios of the predictive variables for the different spatial risk models

Table S2. Risk ratio and 95% credible intervals of the predictive variables in each of the models. The coefficient for the variable *duck* should be interpreted at zero *other poultry* houses, as there is an interaction between *duck* and *other poultry*. Similarly, *other poultry* should be interpreted at zero *duck* houses.

| **Variable** | **2020-2021** | **2021-2022** | **2022-2023** |
|---|---|---|---|
| Number of duck houses | 1.06 [1.05, 1.08] | 1.11 [1.10, 1.13] | 1.13 [1.09, 1.17] |
| Number of other poultry houses | 1.02 [0.99, 1.04] | 1.05 [1.04, 1.07] | 1.07 [1.05, 1.10] |
| Proportion of surface covered by water | 1.02 [0.0007, 622.00] | 0.04 [0.002, 0.76] | 0.61 [0.003, 121.00] |
| Proportion of farms with low competence species | 0.23 [0.09, 0.52] | 0.15 [0.08, 0.27] | 0.59 [0.19, 1.93] |

### Dilution effect for the model 2020-2021

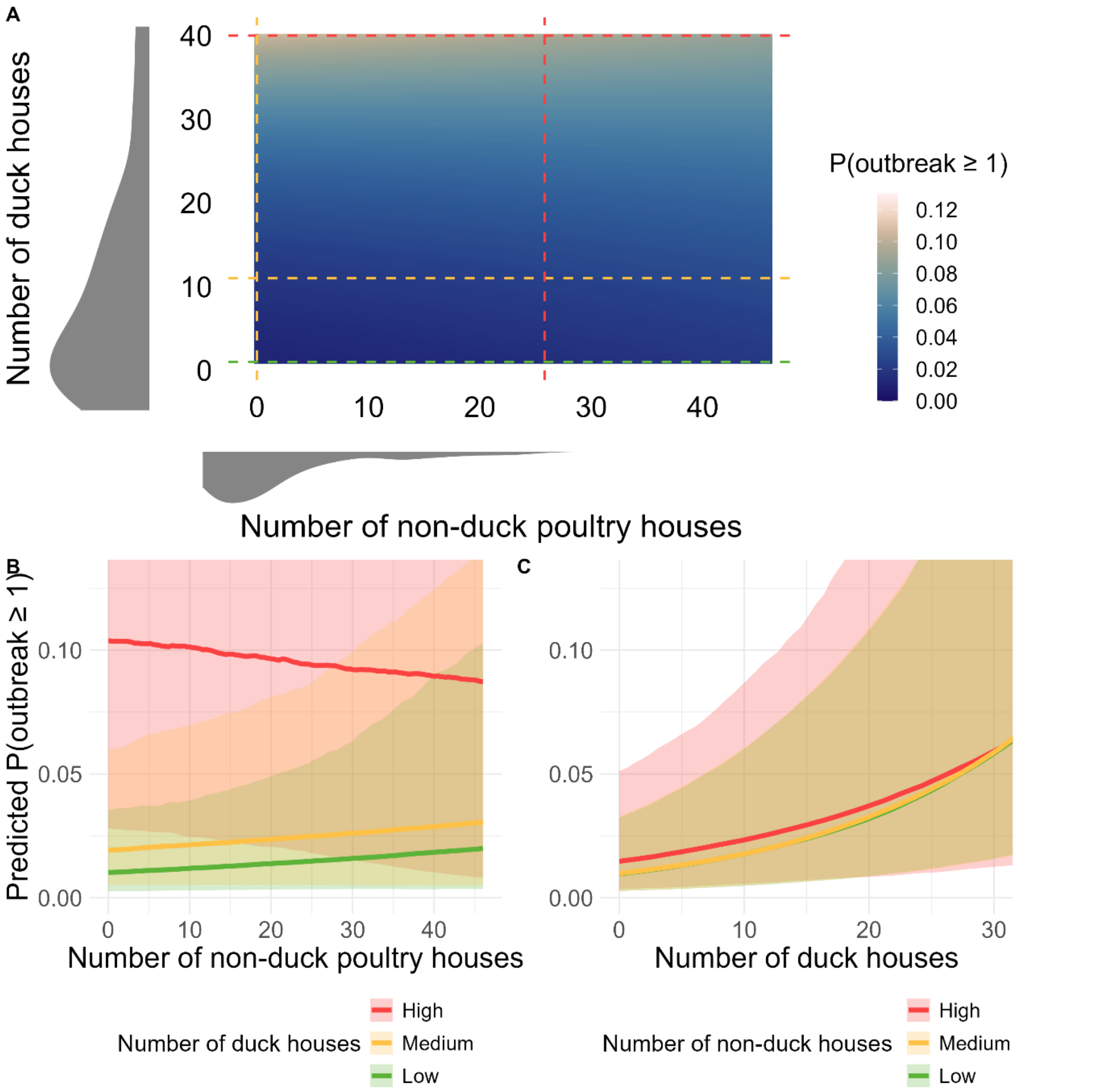


Figure S6. A: Interaction effect between the number of duck and non-duck poultry houses, for the model 2020-2021. The probability of having at least one reported outbreak in the spatial unit as a function of the number of duck houses and number of non-duck poultry houses, calculated for other variables at their average values, without spatial component. The three levels of figure B are represented as horizontal lines, and the three levels of figure C as vertical lines. B: Probability of having at least one reported outbreak in the spatial unit as a function of other poultry house number, for three levels of numbers of duck houses (2.5th, 50th and 97.5th percentile), other variables set at their average values, calculated without spatial component. C: Probability of having at least one reported outbreak in the spatial unit as a function of duck house number, for three levels of numbers of non-duck poultry houses (2.5th, 50th and 97.5th percentile), other variables set at their average values, calculated without spatial component.

### Dilution effect for the model 2022-2023

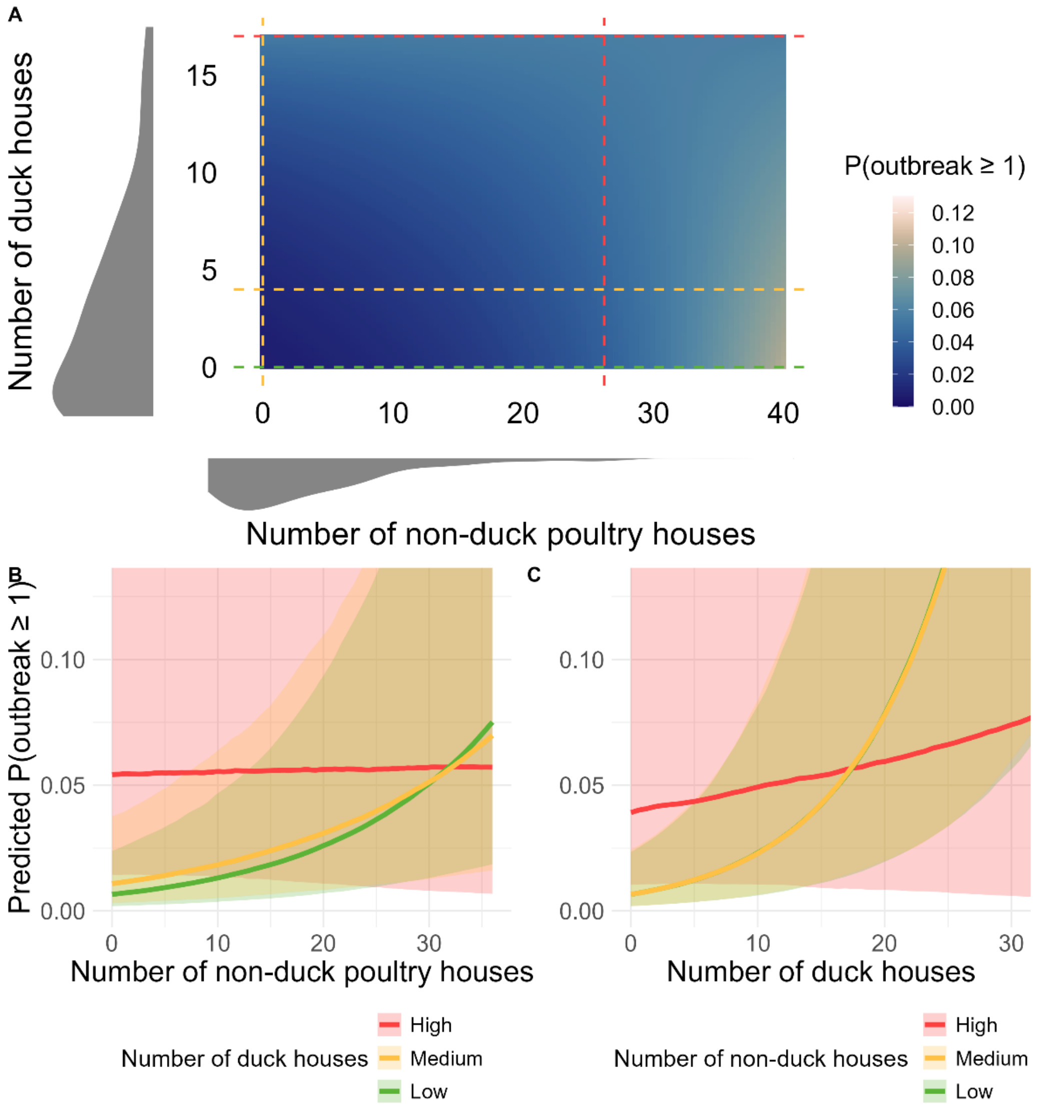


Figure S7. A: Interaction effect between the number of duck and non-duck poultry houses, for the model 2022-2023. The probability of having at least one reported outbreak in the spatial unit as a function of the number of duck houses and number of non-duck poultry houses, calculated for other variables at their average values, without spatial component. The three levels of figure B are represented as horizontal lines, and the three levels of figure C as vertical lines. B: Probability of having at least one reported outbreak in the spatial unit as a function of other poultry house number, for three levels of numbers of duck houses (2.5th, 50th and 97.5th percentile), other variables set at their average values, calculated without spatial component. C: Probability of having at least one reported outbreak in the spatial unit as a function of duck house number, for three levels of numbers of non-duck poultry houses (2.5th, 50th and 97.5th percentile), other variables set at their average values, calculated without spatial component.
